# Biophysical characterization and analysis of a mesophilic chorismate mutase from *B. pumilus*

**DOI:** 10.1101/2023.04.20.537678

**Authors:** Ryan Scott Wilkins, Bjarte Aarmo Lund, Geir Villy Isaksen, Johan Åqvist, Bjørn Olav Brandsdal

**Affiliations:** Hylleraas Centre for Quantum Molecular Sciences, Department of Chemistry, University of Tromsø, N9037 Tromsø, Norway; Department of Cell and Molecular Biology, Uppsala University, Box 596, 75124 Uppsala, Sweden

## Abstract

Chorismate mutases have extensively been used as computational benchmarking systems for enzyme catalysis, yet the roles entropy and enthalpy play in catalysis are still not fully understood. Thus, it is important to better understand these enzymes for potential research or industrial applications. Here, we report the first crystal structure and kinetic characterization of a chorismate mutase from *Bacillus pumilus*. This enzyme exhibits a high degree of similarity to a known mesophilic chorismate mutase from *Bacillus subtilis*. Using this crystal structure, we further employ EVB/MD simulations to construct Arrhenius plots, allowing us to extract thermodynamic activation parameters. Overall, this study provides new insights into the structural and functional features of the *B. pumilus* chorismate mutase and highlights its potential as a valuable enzyme for biocatalytic and biotechnological applications.

## Introduction

In order to better understand the mechanisms behind enzyme catalysis, it is important to know the role the activation enthalpy-entropy balance plays in the reaction. The current hypothesis regarding entropic contributions to enzyme activity proposed by Åqvist *et al*. is that the once-popular Circe effect is an oversimplification^*1*^– activation entropies should not solely include contributions from a loss of translational and rotational entropies upon substrate binding, but also need to account for entropic contributions from the reorganization of the surrounding enzyme and solvent. A common model system used in understanding the balance of these activation parameters in computational chemistry is the chorismate mutase enzyme. ^*2-5*^ Chorismate mutase enzymes are found in bacteria, fungi, and higher plants, where they play a role in the Shikimate pathway. The enzyme is responsible for the catalysis of the pericyclic rearrangement of chorismate to prephenate, which proceeds through a chair-like transition state as seen in Figure (?), which serves as a precursor for the biosynthesis of benzyl-containing amino acids phenylalanine and tyrosine.^*6*^ Further understanding of the roles activation enthalpy and entropy play in enzyme catalysis serves as groundwork for fine-tuning catalytic rates and enzyme stability for the rational design of enzymes to be used in industrial and scientific contexts.

Chorismate mutase enzymes are a common model system to study entropic and enthalpic contributions to enzymatic reactions and are prevalent as a benchmark system for QM/MM calculations. Despite this, there is still debate over how important entropic effects are in the chorismate mutase reaction. Early studies by Gorisch indicate that the chorismate mutase reaction in *S. aurofaciens* is primarily entropy driven. ^*7*^ However later studies on *E. coli* and *B. subtilis* show a significant entropic contribution to the free energy but they are still enthalpically driven reactions. Hur and Bruice originally consider an electrostatic stabilization of the transition state as the C_1_-C_9_ distance decreases the formation of a near-attack conformer (NAC), contributing to the catalytic effect of the enzyme. ^*8*^ On the other hand, Warshel *et al*. later argue that this confirmation is likely a result of electrostatic stabilization, rather than the formation of a NAC. ^*2*^

Preorganization of the substrate into a compressed conformer appears to be a key component underlying the enthalpy-entropy balance in chorismate mutase enzymes, thus understanding the mechanisms leading to this conformer plays a vital role in understanding the role enthalpy and entropy play in the reaction. There has been some work done in identifying key active site residues contributing to the formation of this compressed conformer. In BsCM, substitution of the positively charged Arg90 for a neutral Citrulline results in a significant reduction of *k*_cat_, while only moderately affecting *K*_m_, supporting electrostatic stabilization as a key mechanism in the catalytic reaction. ^*9, 10*^ In the promiscuous enzyme PchB, a lower *k*_cat_ in comparison to monofunctional BsCM has been suggested to be the result of a less restrictive active site. This is supported by an Ala38Ile mutation in *E. coli* CM, ^*11*^ increasing electrostatic stabilization and resulting in a significant increase in *k*_cat_. Additionally, Val35Ile and Val35Ala mutations in EcCM result in an increase in *k*_cat_ of 1.5 times and decrease in *k*_cat_ by about 2 times, respectively. This suggests that chorismate mutase activity can be increased by favoring more restrictive active sites, contributing to higher substrate preorganization and reducing the entropic penalty paid in the reaction. ^*4*^

In this study, we present a new crystal structure for *B. pumilus* chorismate mutase (uniprot A8FEK3, BpCM) obtained from X-ray diffraction crystallography along with thermodynamic activation parameters obtained from constructing an Arrhenius plot from UV-Vis spectroscopy enzyme kinetics experiments. As a full understanding of the roles entropy and enthalpy play in BpCM requires intimate knowledge of atomistic-level details, we further employ computer calculations to elucidate the microscopic details of the enzyme-catalyzed reaction. For this, we construct Arrhenius plots using the empirical valence bond method (EVB) in tandem with molecular dynamics (MD) simulations to obtain free energy calculations of this reaction. Compared to *ab initio* QM/MM methods, EVB maintains a high level of accuracy while remaining less computationally intensive. ^*12-14*^Combining this with molecular dynamics allows us to obtain trajectories of the enzyme and substrate during the reaction. To construct the Arrhenius plot from simulation data, we performed simulations at 7 evenly spaced temperatures ranging from 283-313K on a microsecond-timescale.

## Methods

### Protein production and expression

The gene encoding the *Bacillus pumilus* AroH-type Chorismate mutase (uniprot A8FEK3, BpCM) with a dual C-terminal stop codon (TAATAA) was codon optimized for expression in *E. coli*, synthesized and subcloned into pET22b by GenScript Biotech (Netherlands) B.V. The plasmid was transformed into *E. coli* BL21 (DE3) Star (Invitrogen) using standard heat shock protocol. Protein was produced in shake flasks using ZYP-5052 auto induction media with 50 μg/mL Ampicillin. Cells were grown at 37°C for 4-5 hours, before the temperature was reduced to 20 °C for overnight expression. Cells were harvested by centrifugation, 6000 rpm for 30 minutes. Cells were disrupted by sonication, for 15 minutes active at 40 % amplitude, 3/6 s pulses on/off, 12 °C maximum temperature. The cell lysate was clarified by centrifugation, 20000 rpm for 45 minutes. The clarified cell lysate was loaded on a HisTrap FF 1 mL column equilibrated with 50 mM HEPES pH 7.5 with 500 mM NaCl. Contaminating proteins were eluted by a stepwise elution of 5 and 10 % buffer B containing 500 mM Imidazole and 500 mM NaCl pH 7.5. The protein was eluted by 75 % buffer B. For polishing the concentrated fractions were loaded on a Superdex 75 26/600 column equilibrated with 50 mM potassium phosphate Ph 7.5. The main peak was concentrated to 1 mM concentration (as determined by UV280 measurements with the calculated extinction coefficient) and flash-frozen in liquid nitrogen for storage at -20°C. SEC-MALS was performed with a Shimadzu UFLC system with a Wyatt dawn 8+ detector using Shodex HPLC lb804/lb805 columns with 0.1M NaNO_3_ with 20 mg/L NaN_3_ as running buffer at 50 °C using a protein concentration of 0.65 mg/mL.

### Crystallization

From a set of four commercial crystallization screens from Molecular Dimensions multiple hits were identified for BpCM when screened at 1 mM. For optimization simple gradients of PEG-concentrations were used and diffraction-quality crystals were obtained with 25 % PEG 1500 and 0.1 M MES pH 6.5. Diffraction data was collected using the MX14.2 beamline at the BESSY II electron storage ring at the Helmholtz-Zentrum Berlin für Materialien und Energie Data was processed in XDS.

Molecular replacement was performed with Phaser ^*15*^ using the trimer of *Bacillus subtilis* chorismate mutase 3ZO8 ^*9*^ as the search model. The solution was improved with Autobuild ^*16*^ before several cycles of semi-manual rebuilding in coot ^*17*^ and automatic refinement in phenix.refine. ^*18*^

### Chorismic acid assay

A 10.5 mM stock solution was used to make dilutions from 500 μM to 16 μM. The concentration of BpCM was kept at 10 nM for the experiments. The concentration of chorismic acid was followed at 274 nm, and concentrations were calculated using an extinction coefficient of 2630 mol^-1^cm^-1^. Measurements were done with 1 cm UV-quartz cuvettes in a qChanger6 Peltier-cooled thermostat and sample-changer with a Cary60 UV/VIS spectrophotometer. The velocities from triplicated measurements were fitted against the substrate concentrations with non-linear regression using GraphPad PRISM 6.07 for Windows (GraphPad Software, La Jolla California USA).

### Differential scanning calorimetry

Using 50 mM potassium phosphate pH 7.5 as the buffer and reference the temperature range of 20-80°C was scanned at a rate of 1°C/min.

#### EVB Simulations

The X-ray crystal structure obtained in this study was used as the starting structure for our EVB simulations. The chorismate substrate configuration was obtained by manually superimposing the BpCM crystal structure with the structure of Arg90Cit BsCM with PDB entry 3ZP4. ^*9*^ Topologies for the system were generated using the EVB/MM program *Q*^*19*^ interfaced with *Qgui*, ^*20*^ using the OPLS-AA/M force field to parameterize the enzyme. ^*21*^ A 38 Å spherical boundary centered on the protein center-of-mass was defined around the enzyme, which was filled with TIP3P^*22*^ water molecules to solvate the system. Before the production runs of EVB/MD simulations, the systems were equilibrated by gradually increasing the temperature in 6 steps from 1 K up until the final system temperature. Equilibration was performed in 1 fs timesteps, with the first 5 equilibrations steps being ran for 10 ps, and the final step being performed for 100 ps. System temperature was controlled via coupling to an external heat bath^*23*^ with a relaxation time of 10 fs for the first 5 equilibration steps and 100 fs for the final equilibration step and the production runs. A multipole expansion method (LRF)^*24*^ was used to calculate longrange electrostatics, and nonbonded interactions were calculated using a 10 Å cut-off. However, reacting fragments in the catalytic center were free to interact with the entire system. The SHAKE^*25*^ algorithm was used to constrain bonds and angles of solvent molecules.

The FEP umbrella sampling^*26, 27*^approach was used to calculate EVB free energy profiles. Calculations were divided into 51 discrete FEP windows with a spacing of 0.02 between chorismate (*λ* = 0) and prephenate (*λ* = 1). The EVB Hamiltonian was parameterized using the uncatalyzed conversion of chorismate to prephenate in our previous work. To obtain thermodynamic activation parameters for the enzyme, the EVB/Arrhenius^*29, 30*^ approach was carried out at 7 evenly spaced temperatures from 283-313 K. At each temperature, 100 independent replicas were carried out for a total of 357 ns of simulated trajectories.

#### Molecular Dynamics Simulations

The initial solvated structure for molecular dynamics (MD) simulations was the same structure used in the setup of EVB simulations. Equilibration, electrostatics, non-bonded interactions, and force fields were also identical to those used in the EVB simulations. However, the active site was treated entirely with molecular mechanics. For production runs, 20 independent runs at 3.5 ns of simulation time per run were performed, for a total of 70 ns of simulation time.

## Results and Discussion

### Protein characterization

BpCM was expressed with reasonable yields of 5-10 mg per liter of culture and appeared to be stable. From differential scanning calorimetry a single transition was observed with a T_m_ of 65.5°C compared to 58.3°C for BsCM. No refolding was observed after cooling. SEC-MALS confirmed that BpCM is a trimer in solution.

### Protein structure

The overall structure of BpCM is nearly identical to BsCM, with a RMSD value of 0.28 Å over 717 atoms in the monomers. The RMSD value is surprisingly low as the sequence identity is only 69%, which is on the order of two randomly selected structures of the same protein with 100% identity. However, the trimeric quaternary structure of these chorismate mutases may enforce a certain structure in order to maintain the binding interfaces, forcing any mutations to be compliant within the structural constraints. Calculating the electrostatics using the APBS^*31*^ approach indicates that the BpCM chorismate binding site is more positive than its BsCM homolog (Figure 1). Sitemap^*32, 33*^ calculations of the three binding pockets of the trimers also show that the ratio of hydrogen bond donors to acceptors are higher for the BpCM. Meanwhile BpCM has a lower calculated pI than BsCM (5.33 compared to 5.6) and fewer positively charged amino acids (14 compared to 16) showing that this is a local effect.

**Figure 1:**
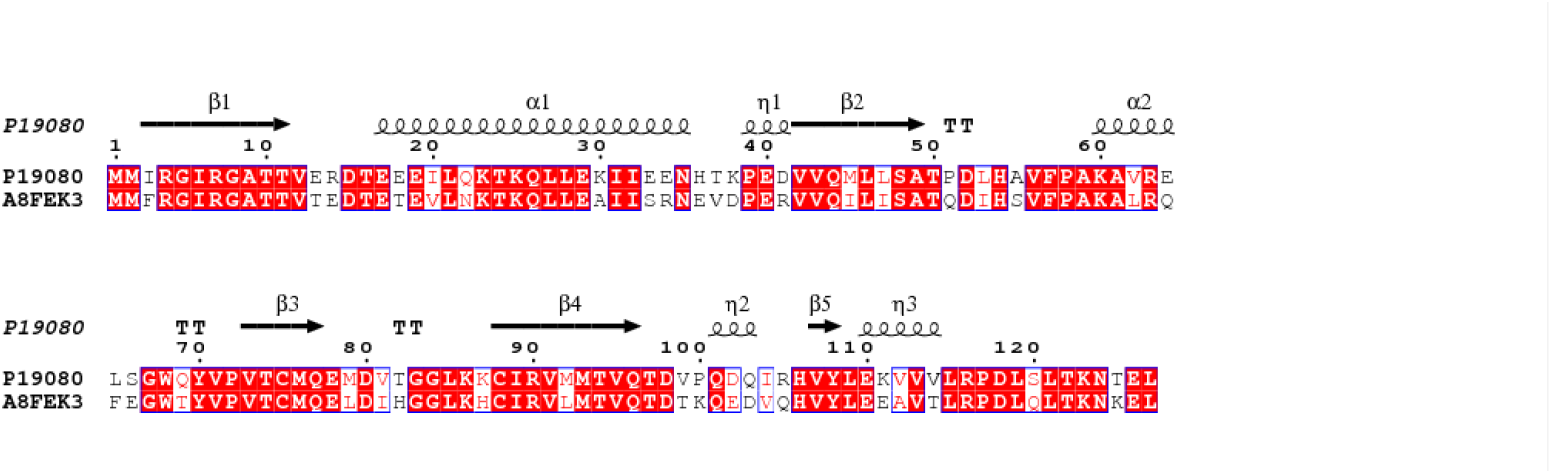
Sequence alignment of BsCM (P19080) and BpCM (A8FEK3) shows an identity of 69%. Furthermore, the similarity between the sequences is 93%.

**Figure 2:**
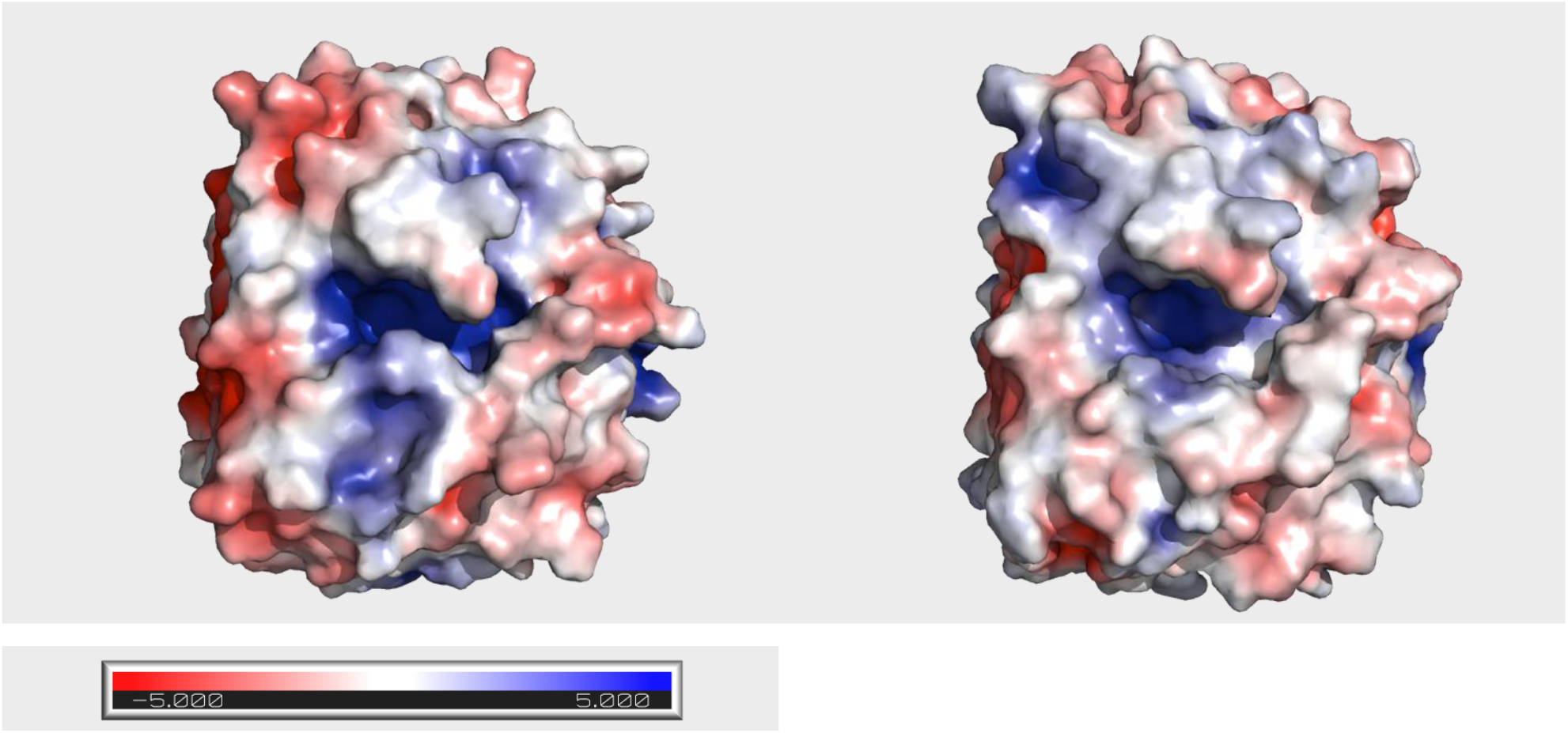
Electrostatics of BpCM and BsCM trimers indicate that the chorismate binding pocket is more positively charged in BpCM than in BsCM. Units given in kT/e

**Figure 3:**
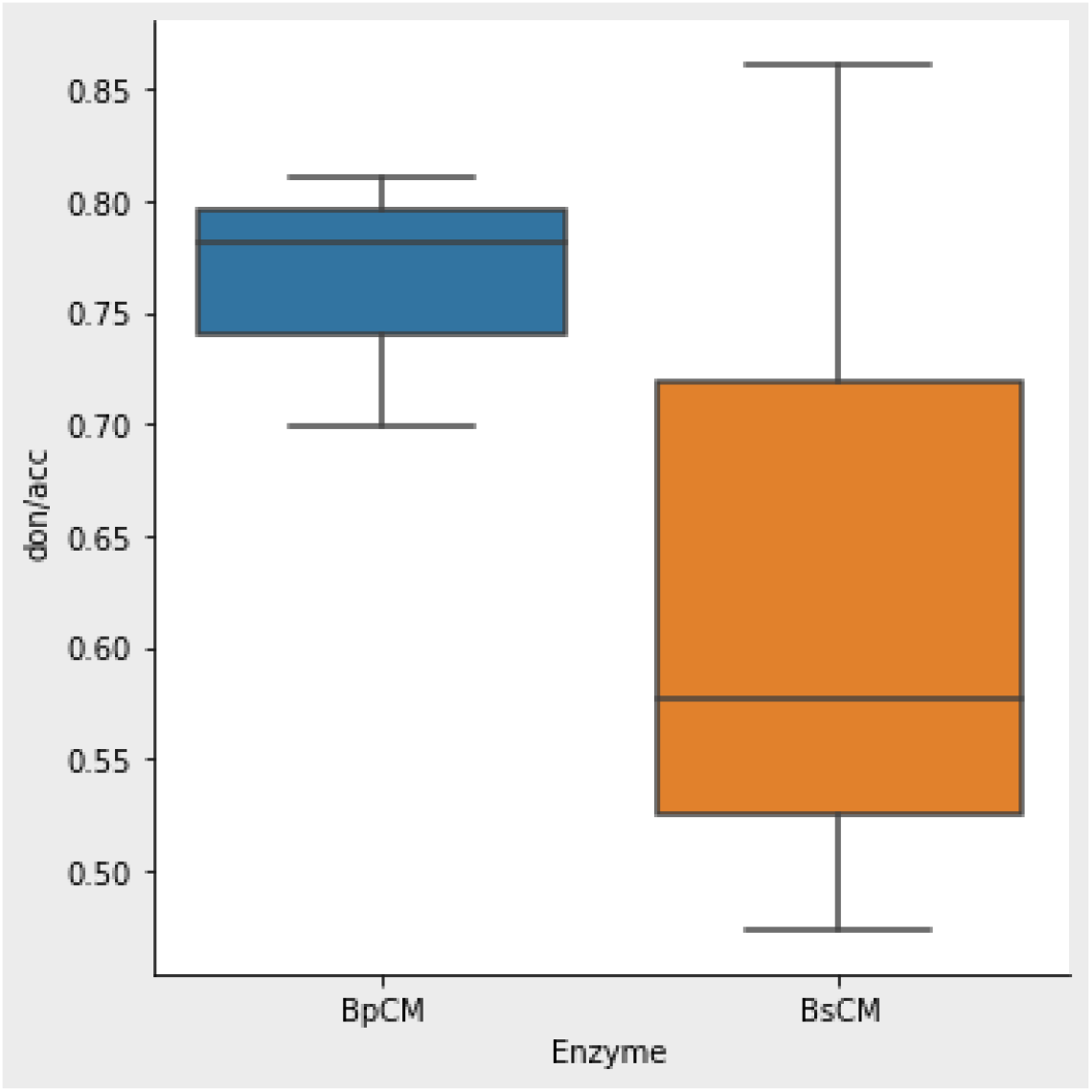
Comparison of binding pockets using SiteMap shows that the donor/acceptor ratio is higher for BpCM than for BsCM

### Activation Parameters of B. pumilus Chorismate Mutase

Using the same intrinsic gas phase energy gap Δα and off-diagonal coupling H_12_ calibrated in our previous work on BsCM, we performed EVB simulations on BpCM at 7 different temperatures ranging from 283-313 K, from which an Arrhenius plot was constructed. Fitting Δ*G*^‡^/*T* vs. 1/*T* using a linear regression, we obtained thermodynamic activation parameters of Δ*H*^‡^ = 12.9 ± 0.5 kcal mol^-1^ and *T*Δ*S*^‡^ = *-*2.1 ± 0.5 kcal mol^-1^. An Arrhenius plot and similar regression of experimental results, obtained by fitting ln(*k*_cat_) versus temperature values obtained from UV-Vis spectroscopy in a chorismic acid assay, yielded activation parameters of Δ*H*^‡^ = 13.1 ± 0. kcal mol^-1^ and *T*Δ*S*^‡^ = *-*2.1 ± 0.5 kcal mol^-1^. The thermodynamic activation parameters predicted from the Arrhenius plot approach applied to both simulation and experimental results are in excellent agreement with each other.

Compared to the related chorismate mutase enzyme from *B. subtilis* (BsCM), which shares a 69% sequence identity and 93% similarity to BpCM, the thermodynamic activation parameters also indicate that like BsCM, BpCM is likely a mesophilic enzyme, despite other enzymes from *B. pumilus* exhibiting psychrophilic behavior. ^*35, 36*^ The activation free energy for BpCM is less than 1 kcal mol^-1^ lower than reported values for BsCM, both experimentally and computationally. The activation enthalpies of both enzymes are the same within reported errors, thus the difference in **Δ**G^‡^ is likely a result of BpCM having a lower *T***Δ**S^‡^ of about 0.6 kcal mol^-1^ at 298 K.

### Structural Analysis of the Catalytic Effect of B. pumilus Chorismate Mutase

To further understand the mechanisms contributing to the catalytic effect in BpCM, we compare it to the structure to of BsCM, which has been well documented. According to our analysis of the thermodynamic activation parameters of the two enzymes, the differences between the two appears to mostly due to entropic effects, as the activation enthalpies of the two enzymes are remarkably similar.

An analysis of the active sites using both a 3D snapshot of the active site during the transition state (Figure 4) and a 2D ligand interaction diagram (Figure 5) generated in Maestro (Schrödinger Release 2019-3) ^*37*^ indicates a highly similar active site between the two enzymes, with identical hydrogen bonding between active site residues in both enzymes and the substrate, with the exception of the presence of an additional hydrogen bond in the case of BpCM, between Cys75 and the hydroxyl group attached to the C_4_ atom on the substrate. However, an analysis of trajectories generated from the molecular dynamics portion of our EVB simulations indicates a distance of 3.2 ± 0.2 Å between the main chain amide proton in Cys75 of BsCM, compared to a distance of 2.9 ± 0.2 Å with the corresponding Cys75 in BpCM (Figure 6). This is indicative that there is also hydrogen bonding present between this residue and the substrate in BsCM, however it lies outside of Maestro ‘s distance cut-off. Hydrogen bonding between these residues and the C_4_-OH are responsible for an elongation of the C_5_-O_7_ bond in chorismate, leading to a stabilization effect in the transition state of the substrate.^*38*^ Mutagenesis studies where Glu78 was mutated to residues disrupting this hydrogen bonding indicate a reduction in the catalytic rate of BsCM, supporting the role for these residues in the catalytic effect of the enzymes.

**Figure 4:**
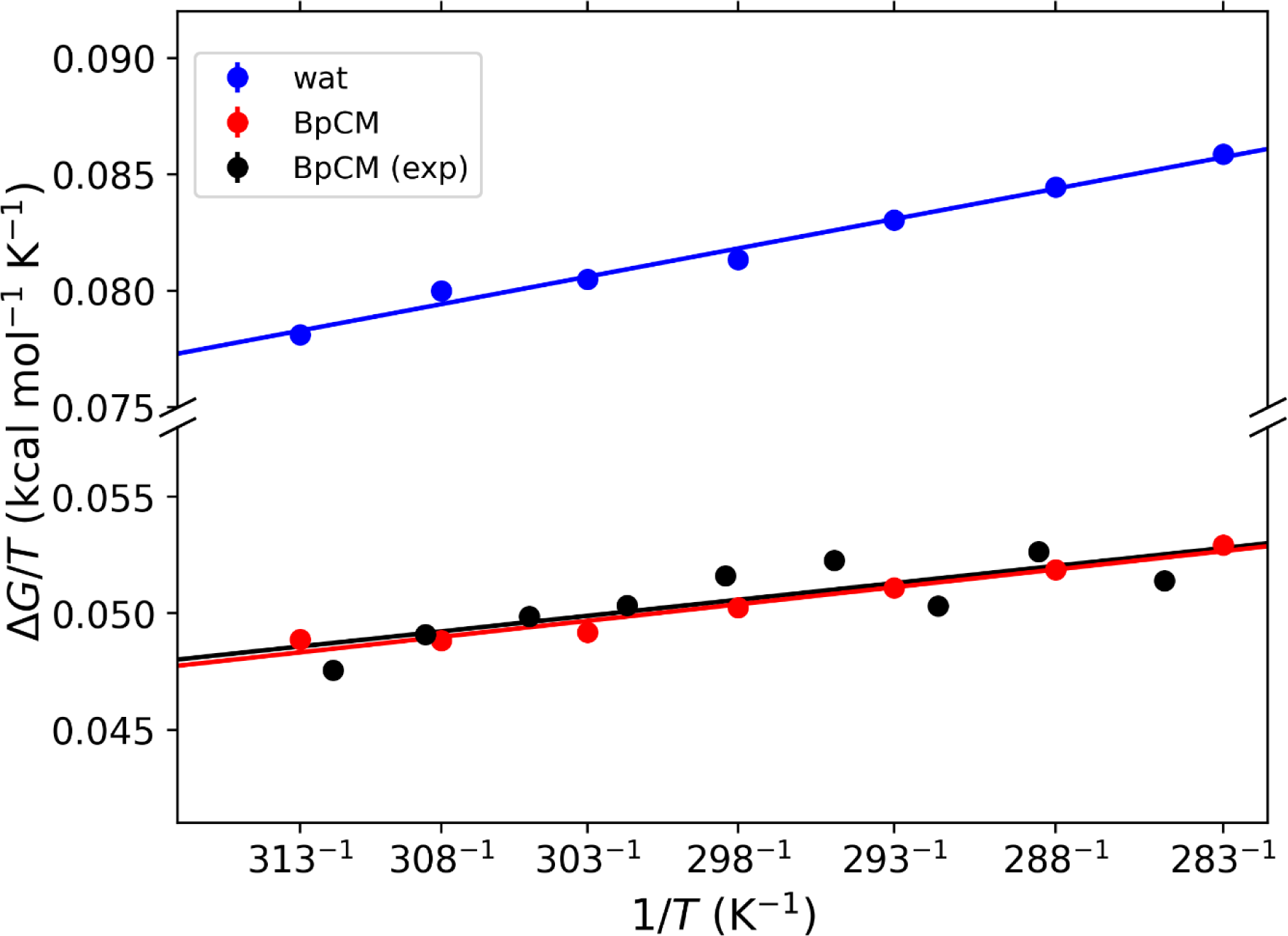
Arrhenius plots from 283-313 K comparing spectrophotometry (black) and EVB results for BpCM to EVB results of the reaction in an aqueous environment (blue).

**Figure 4:**
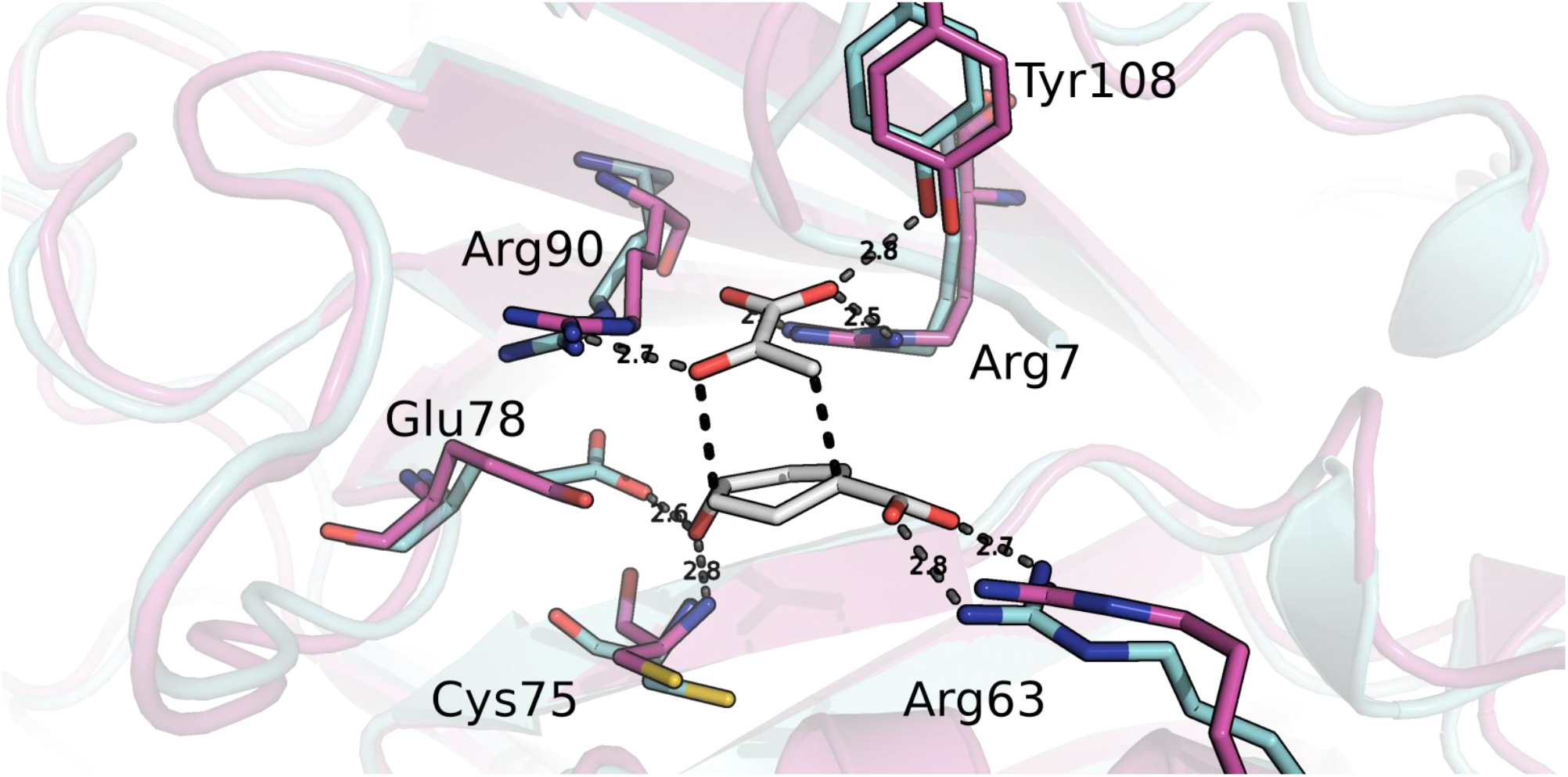
Active site comparison of BsCM (cyan) and BpCM (pink) with the transition state complex (white) obtained from the trajectory of MD simulations showing the highly conserved residues of interest.

**Figure 5:**
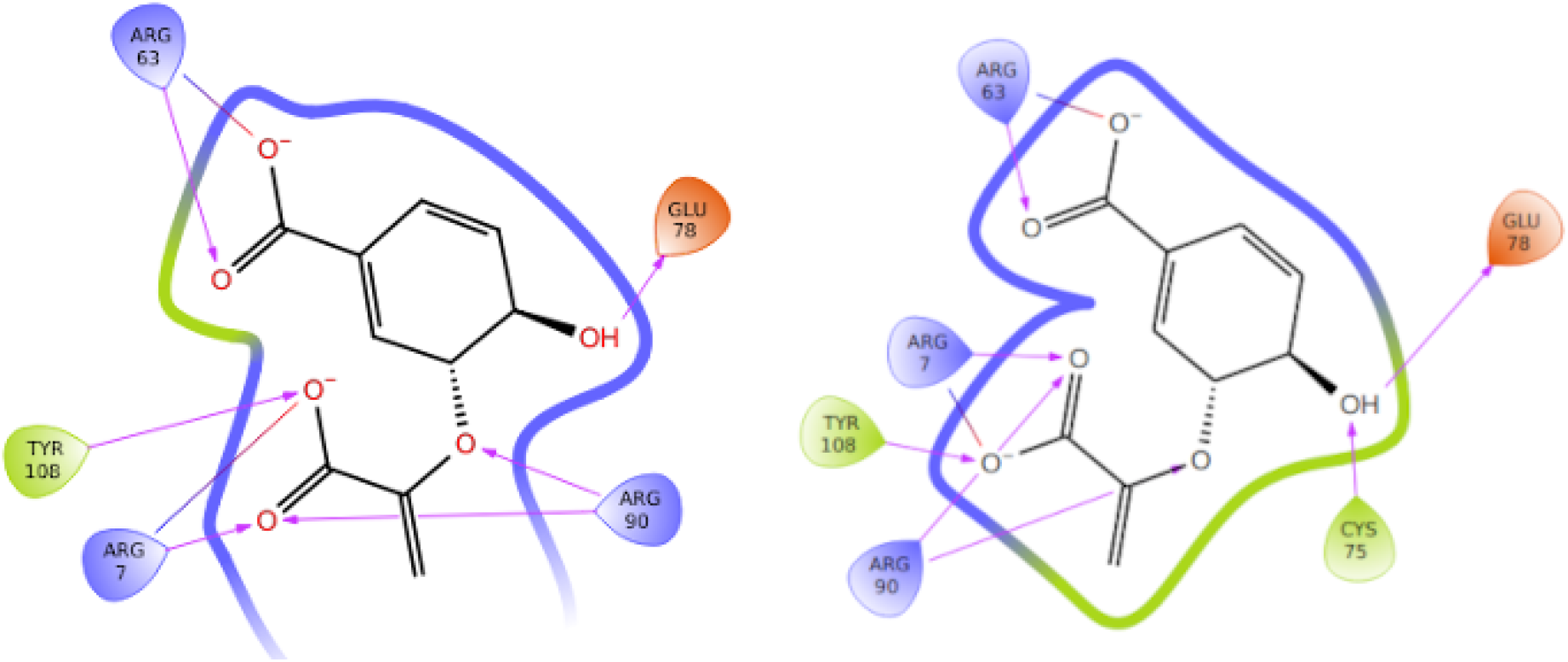
Ligand interaction diagrams of (A) BsCM and (B) BpCM showing a high degree of similarity in the active sites of the two enzymes.

**Figure 6:**
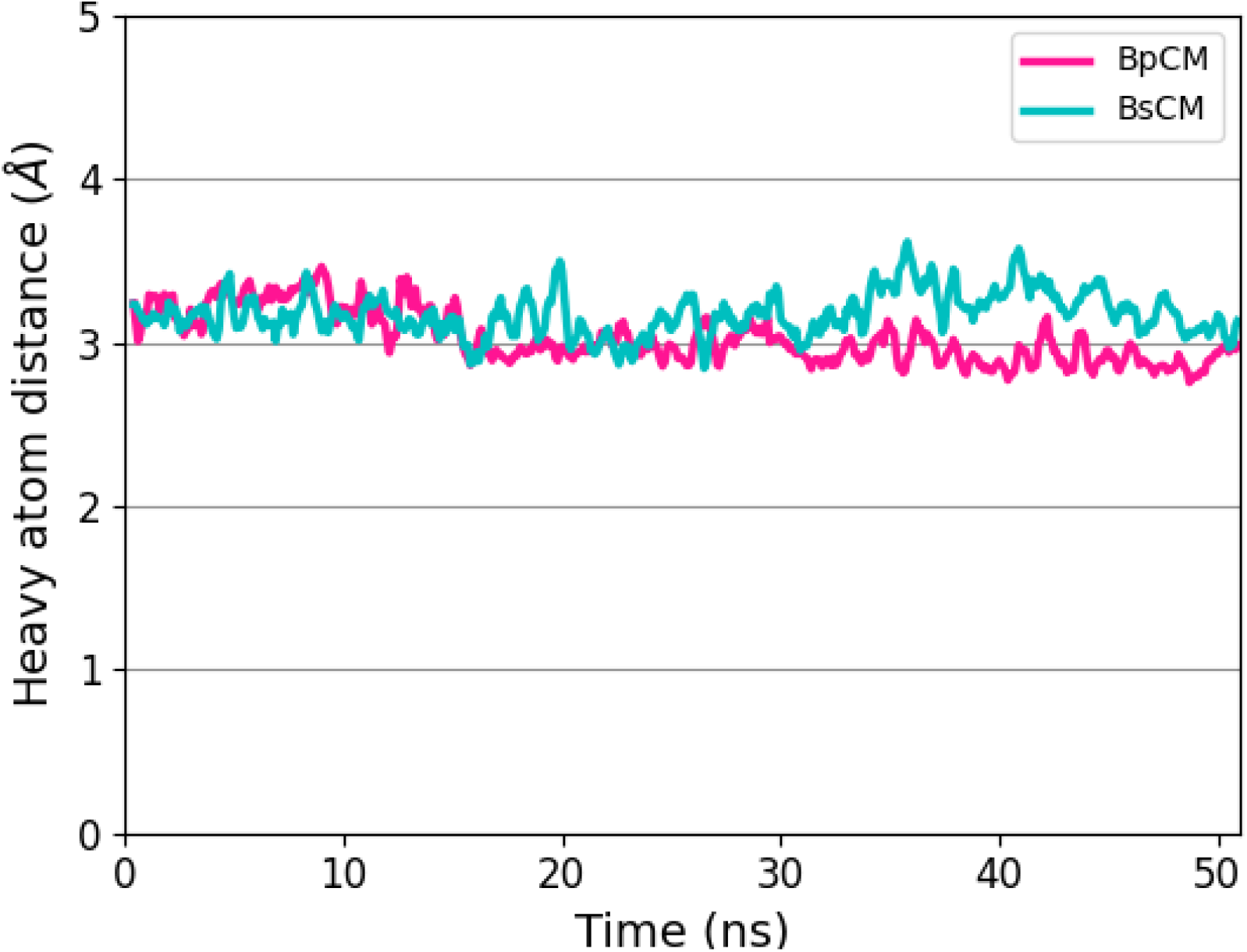
Heavy atom distances between the substrate hydroxyl group on C_4_ and amide protons of Cys75 (BpCM in pink, BsCM in cyan) throughout the course of the chorismate mutase reaction.

**Figure 7:**
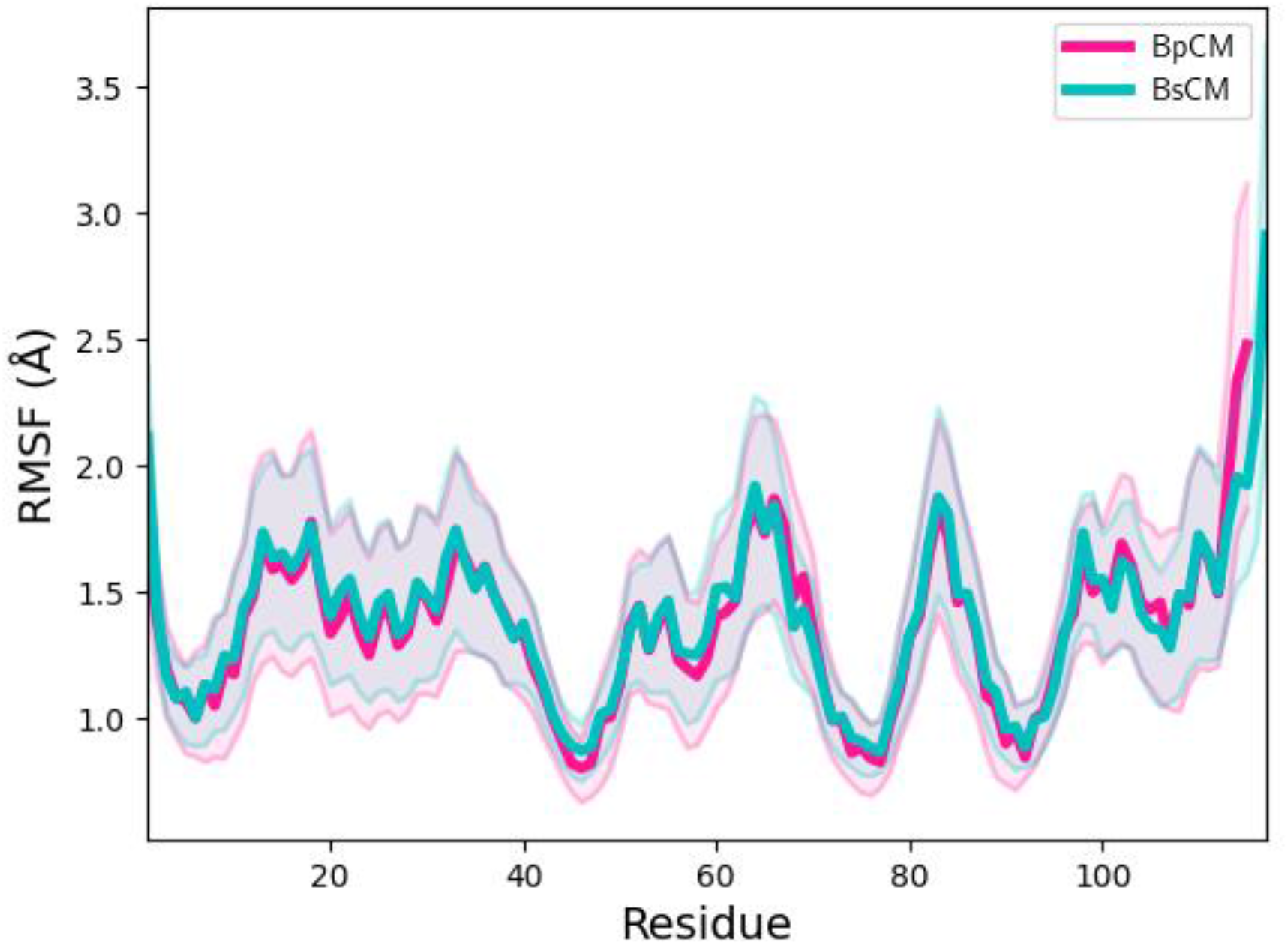
Root mean square fluctuations per residue of each monomer averaged over the N-C_α_-C-O backbone atoms in BpCM (pink) versus BsCM (cyan) with error bands shown in respective colours.

Furthermore, a large negative charge is generated on the ether oxygen (the O_7_ atom, see FigureX) in the transition state by the elongation of the C_5_-O_7_ ether bond. In BpCM, the proximity of Arg90 likely provides a positive charge to counterbalance this effect. This corresponds directly to the Arg90 in BsCM fulfilling the same function due to the high active site similarity. Thus, the presence of these arginines in the active site is likely to help provide a positive charge stabilize a negatively charged region in transition state.

Formation of the C_1_-C_9_ bond is crucial in the formation of prephenate. To form this bond, the pyruvate moiety in the transition state must maintain a proper orientation with respect to the cyclohexadienyl moiety. Mutagenesis studies on a related *E. coli* CM show a 10^4^ reduction in *k*_cat_/*K*_m_ when residues orienting the pyruvate moiety were mutated to prevent proper alignment during the transition state. ^*39*^ To maintain the proper orientation, Arg7 and Tyr108 in BpCM form strong ionic and hydrogen bonds with the pyruvate moiety in the transition state. Similarly, in BsCM these bonds to the pyruvate moiety are made with the same Arg7 and Tyr108.

An analysis of residue fluctuations in both enzymes was calculated from a total 35ns of molecular dynamics trajectories over 10 independent runs. Due to the high sequence identity and similarity between the two enzymes, the RMSF profiles are remarkably similar, with only slight deviations corresponding to residues found mostly in surface loop regions. An exception can be found in residue 45, found in the middle of the β-sheets forming the internal core of the enzymes, in which a polar Met45 in BsCM is found in pocket formed otherwise by hydrophobic residues. In contrast, the corresponding residue in BpCM is Ile45, which is likely to be more stable in this hydrophobic pocket.

To further understand the internal energetics of the reaction, we can perform a breakdown into bonded and nonbonded contributions (Table 2), as seen in our previous work (cite JCTC paper here when available). The total internal energy of the reacting region in BsCM is slightly more favorable than that of the BpCM, likely due to slightly less favorable nonbonded interactions in BpCM despite slightly more favorable bonded interactions. Analyzing the contributions to 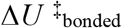 shows more favorable torsional and improper contributions to the internal energy, while contributions from bond and angle interactions are approximately same in both BsCM and BpCM. A similar breakdown of the nonbonded interactions shows that while van der Waals contributions to the nonbonded internal energy are also similar between the two enzymes, the active site of BsCM provides a slightly more favorable electrostatic environment.

**Table 1:** Thermodynamic activation parameters for the chorismate mutase (CM) reaction (chorismate → prephenate) in water (uncatalyzed) and catalyzed by chorismate mutase enzymes from *B. pumilus* (BpCM) and *B. subtilis* (BsCM).

| | $\Delta G^\ddagger$<br>(kcal/mol) | $\Delta H^\ddagger$ (kcal/mol) | $T\Delta S^\ddagger$ (cal mol<sup>-1</sup> K<sup>-1</sup>) |
| --- | --- | --- | --- |
| **Uncatalyzed** |  |  |  |
| Experimental<sup>a</sup> | 24.5 | 20.7 | -3.8 |
| EVB<sup>c</sup> | 24.4 ± 0.6 | 20.9 ± 0.4 | -3.4 ± 0.4 |
| **BpCM** |  |  |  |
| Experimental | 15.2 | 13.1 | -2.1 |
| EVB | 15.0 ± 0.7 | 12.9 ± 0.5 | -2.1 ± 0.5 |
| **BsCM** |  |  |  |
| Experimental<sup>b</sup> | 15.4 | 12.7 ± 0.4 | -2.7 ± 0.4 |
| EVB<sup>c</sup> | 15.8 ± 1.2 | 12.9 ± 0.8 | -2.9 ± 0.3 |

<sup>a</sup>Andrews et al.<sup>28</sup> <sup>b</sup>Kast et al.<sup>34</sup> <sup>c</sup>Wilkins et al.

**Table 2:**
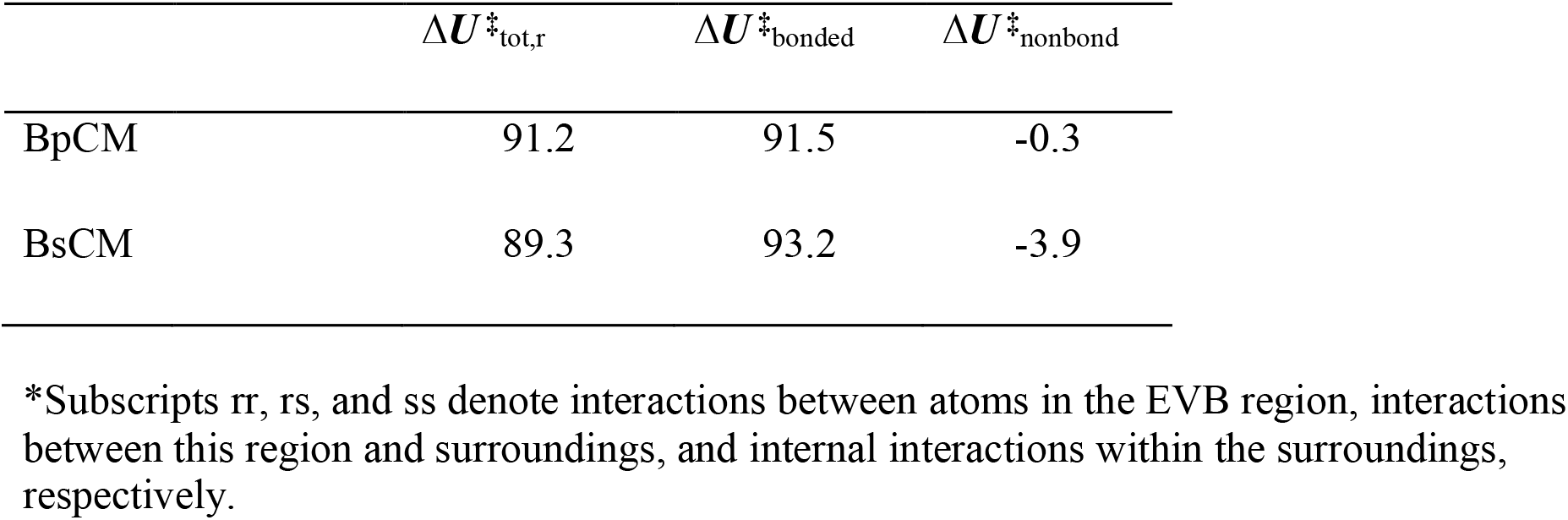
Total potential activation energy within the reacting region and surroundings (in kcal/mol at 298 K) for the chorismate mutase (CM) reaction (chorismate → prephenate) from *B. pumilus* (BpCM) and from *B. subtilis* (BsCM), respectively.

| | $\Delta U^{\ddagger}_{\text{tot,r}}$ | $\Delta U^{\ddagger}_{\text{bonded}}$ | $\Delta U^{\ddagger}_{\text{nonbond}}$ |
| --- | --- | --- | --- |
| BpCM | 91.2 | 91.5 | -0.3 |
| BsCM | 89.3 | 93.2 | -3.9 |

\*Subscripts rr, rs, and ss denote interactions between atoms in the EVB region, interactions between this region and surroundings, and internal interactions within the surroundings, respectively.

## Concluding remarks

We have presented a new crystal structure and accompanying characterization of thermodynamic activation parameters for a new chorismate mutase enzyme from *Bacillus pumilus*. The sequence and structure show high similarity to the previously characterized *B. subtilis* chorismate mutase. Thermodynamic activation parameters obtained from experiment and molecular simulation, along with an accompanying structural analysis, show enthalpic contributions to the reaction pathway are identical, but there exists a small difference in activation entropy between BpCM and BsCM.

Understanding the thermodynamic activation parameters and the underlying structural details is a crucial component leading to rational enzyme design. Understanding the way in which residues in the enzyme contribute to enthalpic and entropic effects in the chorismate mutase reaction establishes BpCM as a possible starting point in the design of psychrophilic or thermophilic chorismate mutase enzymes.

## ACKNOWLEDGMENT

Support from the Norwegian Research Council through a Centre of Excellence and project grant (Grant Nos. 262695 and 274858), the Swedish Research Council (VR) and the Knut and Alice Wallenberg Foundation is gratefully acknowledged.

## References

[1] Kazemi, M., Himo, F., and Aqvist, J. (2016) Enzyme catalysis by entropy without Circe effect, Proc Natl Acad Sci U S A 113, 2406–2411.

[2] Strajbl, M., Shurki, A., Kato, M., and Warshel, A. (2003) Apparent NAC effect in chorismate mutase reflects electrostatic transition state stabilization, J Am Chem Soc 125, 10228–10237.

[3] Lyne, P. D., Mulholland, A. J., and Richards, W. G. (1995) Insights into Chorismate Mutase Catalysis from a Combined QM/MM Simulation of the Enzyme Reaction, Journal of the American Chemical Society 117, 11345–11350.

[4] Marti, S., Andres, J., Moliner, V., Silla, E., Tunon, I., and Bertran, J. (2008) Predicting an improvement of secondary catalytic activity of promiscuous isochorismate pyruvate lyase by computational design, J Am Chem Soc 130, 2894–2895.

[5] Marti, S., Andres, J., Moliner, V., Silla, E., Tunon, I., Bertran, J., and Field, M. J. (2001) A hybrid potential reaction path and free energy study of the chorismate mutase reaction, J Am Chem Soc 123, 1709–1712.

[6] Gibson, F., and Pittard, J. (1968) Pathways of biosynthesis of aromatic amino acids and vitamins and their control in microorganisms, Bacteriol Rev 32, 465–492.

[7] Görisch, H. (1978) A new test for chorismate mutase activity, Analytical Biochemistry 86, 764–768.

[8] Hur, S., and Bruice, T. C. (2003) Enzymes do what is expected (chalcone isomerase versus chorismate mutase), J Am Chem Soc 125, 1472–1473.

[9] Burschowsky, D., van Eerde, A., Okvist, M., Kienhofer, A., Kast, P., Hilvert, D., and Krengel, U. (2014) Electrostatic transition state stabilization rather than reactant destabilization provides the chemical basis for efficient chorismate mutase catalysis, Proc Natl Acad Sci U S A 111, 17516–17521.

[10] Kienhofer, A., Kast, P., and Hilvert, D. (2003) Selective stabilization of the chorismate mutase transition state by a positively charged hydrogen bond donor, J Am Chem Soc 125, 3206–3207.

[11] Lassila, J. K., Keeffe, J. R., Kast, P., and Mayo, S. L. (2007) Exhaustive mutagenesis of six secondary active-site residues in Escherichia coli chorismate mutase shows the importance of hydrophobic side chains and a helix N-capping position for stability and catalysis, Biochemistry 46, 6883–6891.

[12] Åqvist, J., and Warshel, A. (1993) Simulation of enzyme reactions using valence bond force fields and other hybrid quantum/classical approaches, Chemical Reviews 93, 2523–2544.

[13] Kamerlin, S. C., and Warshel, A. (2010) The EVB as a quantitative tool for formulating simulations and analyzing biological and chemical reactions, Faraday Discuss 145, 71–106.

[14] Warshel, A., and Weiss, R. M. (1980) An empirical valence bond approach for comparing reactions in solutions and in enzymes, Journal of the American Chemical Society 102, 6218–6226.

[15] McCoy, A. J., Grosse-Kunstleve, R. W., Adams, P. D., Winn, M. D., Storoni, L. C., and Read, R. J. (2007) Phaser crystallographic software, J Appl Crystallogr 40, 658–674.

[16] Terwilliger, T. C., Grosse-Kunstleve, R. W., Afonine, P. V., Moriarty, N. W., Zwart, P. H., Hung, L. W., Read, R. J., and Adams, P. D. (2008) Iterative model building, structure refinement and density modification with the PHENIX AutoBuild wizard, Acta Crystallogr D Biol Crystallogr 64, 61–69.

[17] Emsley, P., Lohkamp, B., Scott, W. G., and Cowtan, K. (2010) Features and development of Coot, Acta Crystallogr D Biol Crystallogr 66, 486–501.

[18] Afonine, P. V., Grosse-Kunstleve, R. W., Echols, N., Headd, J. J., Moriarty, N. W., Mustyakimov, M., Terwilliger, T. C., Urzhumtsev, A., Zwart, P. H., and Adams, P. D. (2012) Towards automated crystallographic structure refinement with phenix.refine, Acta Crystallogr D Biol Crystallogr 68, 352–367.

[19] Marelius, J., Kolmodin, K., Feierberg, I., and Åqvist, J. (1998) Q: a molecular dynamics program for free energy calculations and empirical valence bond simulations in biomolecular systems, Journal of Molecular Graphics and Modelling 16, 213–225.

[20] Isaksen, G. V., Andberg, T. A., Aqvist, J., and Brandsdal, B. O. (2015) Qgui: A highthroughput interface for automated setup and analysis of free energy calculations and empirical valence bond simulations in biological systems, J Mol Graph Model 60, 15–23.

[21] Robertson, M. J., Qian, Y., Robinson, M. C., Tirado-Rives, J., and Jorgensen, W. L. (2019) Development and Testing of the OPLS-AA/M Force Field for RNA, J Chem Theory Comput 15, 2734–2742.

[22] Jorgensen, W. L., Chandrasekhar, J., Madura, J. D., Impey, R. W., and Klein, M. L. (1983) Comparison of simple potential functions for simulating liquid water, The Journal of Chemical Physics 79, 926–935.

[23] Berendsen, H. J. C., Postma, J. P. M., van Gunsteren, W. F., DiNola, A., and Haak, J. R. (1984) Molecular dynamics with coupling to an external bath, The Journal of Chemical Physics 81, 3684–3690.

[24] Lee, F. S., and Warshel, A. (1992) A local reaction field method for fast evaluation of long-range electrostatic interactions in molecular simulations, The Journal of Chemical Physics 97, 3100–3107.

[25] Ryckaert, J.-P., Ciccotti, G., and Berendsen, H. J. C. (1977) Numerical integration of the cartesian equations of motion of a system with constraints: molecular dynamics of nalkanes, Journal of Computational Physics 23, 327–341.

[26] Warshel, A. (1991) Computer modelling of chemical reactions in enzymes and solutions.

[27] Hwang, J. K., King, G., Creighton, S., and Warshel, A. (1988) Simulation of free energy relationships and dynamics of SN2 reactions in aqueous solution, Journal of the American Chemical Society 110, 5297–5311.

[28] Andrews, P. R., Smith, G. D., and Young, I. G. (1973) Transition-state stabilization and enzymic catalysis. Kinetic and molecular orbital studies of the rearrangement of chorismate to prephenate, Biochemistry 12, 3492–3498.

[29] Isaksen, G. V., Aqvist, J., and Brandsdal, B. O. (2014) Protein surface softness is the origin of enzyme cold-adaptation of trypsin, PLoS Comput Biol 10, e1003813.

[30] Kazemi, M., and Aqvist, J. (2015) Chemical reaction mechanisms in solution from brute force computational Arrhenius plots, Nat Commun 6, 7293.

[31] Jurrus, E., Engel, D., Star, K., Monson, K., Brandi, J., Felberg, L. E., Brookes, D. H., Wilson, L., Chen, J., Liles, K., Chun, M., Li, P., Gohara, D. W., Dolinsky, T., Konecny, R., Koes, D. R., Nielsen, J. E., Head-Gordon, T., Geng, W., Krasny, R., Wei, G. W., Holst, M. J., McCammon, J. A., and Baker, N. A. (2018) Improvements to the APBS biomolecular solvation software suite, Protein Sci 27, 112–128.

[32] Halgren, T. (2007) New method for fast and accurate binding-site identification and analysis, Chem Biol Drug Des 69, 146–148.

[33] Halgren, T. A. (2009) Identifying and characterizing binding sites and assessing druggability, J Chem Inf Model 49, 377–389.

[34] Kast, P., Asif-Ullah, M., and Hilvert, D. (1996) Is chorismate mutase a prototypic entropy trap? - Activation parameters for the Bacillus subtilis enzyme, Tetrahedron Letters 37, 2691–2694.

[35] Ranaghan, K. E., Shchepanovska, D., Bennie, S. J., Lawan, N., Macrae, S. J., Zurek, J., Manby, F. R., and Mulholland, A. J. (2019) Projector-Based Embedding Eliminates Density Functional Dependence for QM/MM Calculations of Reactions in Enzymes and Solution, J Chem Inf Model 59, 2063–2078.

[36] van der Ent, F., Lund, B. A., Svalberg, L., Purg, M., Chukwu, G., Widersten, M., Isaksen, G. V., Brandsdal, B. O., and Aqvist, J. (2022) Structure and Mechanism of a Cold-Adapted Bacterial Lipase, Biochemistry 61, 933–942.

[37] Schrödinger Release 2019-3: Maestro, Schrödinger, LLC.

[38] Worthington, S. E., Roitberg A. E., and Krauss, M. (2001) An MD/QM Study of the Chorismate Mutase-Catalyzed Claisen Rearrangement Reaction, The Journal of Physical Chemistry B 105, 7087–7095.

[39] Liu, D. R., Cload, S. T., Pastor, R. M., and Schultz, P. G. (1996) Analysis of Active Site Residues in Escherichia coli Chorismate Mutase by Site-Directed Mutagenesis, Journal of the American Chemical Society 118, 1789–1790.

